# Automatic segmentation of dentate nuclei for microstructure assessment: example of application to temporal lobe epilepsy patients

**DOI:** 10.1101/2020.07.29.222430

**Authors:** Marta Gaviraghi, Giovanni Savini, Gloria Castellazzi, Fulvia Palesi, Nicolò Rolandi, Simone Sacco, Anna Pichiecchio, Valeria Mariani, Elena Tartara, Laura Tassi, Paolo Vitali, Egidio D’Angelo, Claudia A.M. Gandini Wheeler-Kingshott

**Affiliations:** Department of Electrical, Computer and Biomedical Engineering, University of Pavia, Pavia, Italy; Neuroradiology Unit, Brain MRI 3T Research Center, IRCCS Mondino Foundation, Pavia, Italy; Queen Square MS Centre, Department of Neuroinflammation, UCL Queen Square Institute of Neurology, Faculty of Brain Sciences, University College London, London, United Kingdom; Brain Connectivity Center (BCC), IRCCS Mondino Foundation, Pavia, Italy; Department of Brain and Behavioral Sciences, University of Pavia, Pavia, Italy; UCSF Weill Institute for Neurosciences, Department of Neurology, University of California, San Francisco; Department of Clinical Surgical Diagnostic and Pediatric Sciences, University of Pavia, Pavia, Italy; Neuroradiology Unit, IRCCS Mondino Foundation, Pavia, Italy; “C. Munari” Centre for Epilepsy Surgery, Grande Ospedale Metropolitano Niguarda, Milan, Italy; Italian National Research Council (CNR), Institute of Neuroscience, Parma, Italy; Epilepsy Centre, IRCCS Mondino Foundation, Pavia, Italy

## Abstract

Dentate nuclei (DNs) segmentation is helpful for assessing their potential involvement in neurological diseases. Once DNs have been segmented, it becomes possible to investigate whether DNs they are microstructurally affected, through analysis of quantitative MRI parameters, such as the ones derived from diffusion weighted imaging (DWI). This study, therefore, aimed to develop a fully automated segmentation method using the non-DWI (b0) images from a DWI dataset to obtain DN masks inherently registered with parameter maps.

Three different automatic methods were applied to healthy subjects in order to segment the DNs: registration to SUIT (a spatially unbiased atlas template of the cerebellum and brainstem), OPAL (Optimized Patch Match for Label fusion) and CNN (Convolutional Neural Network). DNs manual segmentation was considered the gold standard. Results show that the segmentation obtained with SUIT has an average Dice Similarity Coefficient (DSC) of 0.4907±0.0793 between the automatic SUIT masks and the gold standard. A comparison with manual masks was also performed for OPAL (DSC = 0.7624 ± 0.1786) and CNN (DSC = 0.8658 ± 0.0255), showing a better performance when using CNN.

OPAL and CNN were optimised on heathy subjects’ data with high spatial resolution from the Human Connectome Project. The three methods were further used to segment the DNs of a subset of subjects affected by Temporal Lobe Epilepsy (TLE). This subset was derived from a 3T MRI research study which included DWI data acquired with a coarser resolution. In TLE dataset, SUIT performed similarly to using the HCP dataset, with a DSC = 0.4145 ± 0.1023. Using TLE data, OPAL performed worse than using HCP data: after changing the probability threshold the DSC was 0.4522 ± 0.1178.

CNN was able to extract the DNs using the TLE data without need for retraining and with a good DSC = 0.7368 ± 0.0799. Statistical comparison of quantitative parameters derived from DWI analysis, as well as volumes of each DN, revealed altered and lateralised changes in TLE patients compared to healthy controls.

The proposed CNN is therefore a viable option for accurate extraction of DNs from b0 images of DWI data with different resolutions and acquired at different sites.

## 1 Introduction

Cerebellar nuclei (CNs) have a fundamental role in the central nervous system; they are the main output channels of the cerebellum towards the supratentorial brain and the spinal cord (Sure and Culicchia, 2005). The dentate nuclei (DNs) are the CNs with the largest volume (measuring about 2 cm in the anterior-posterior direction and 1 cm in transverse plane and coronal plane) (Cattaneo, 1989) and they are the matter of this study. Histologically, the DNs have the shape of an irregularly pleated grey foil, very thin and with a longitudinal section appearing as a curved line that contains white matter. Its afferent input comes mainly from the cerebellar cortex and its efferent fibers travel via the superior cerebellar peduncle to the contralateral red nucleus and thalamus (Sure and Culicchia, 2005). The DNs are known mainly for their involvement with the sensorimotor system, although recently they have been shown to respond with strong activation also during cognitive tasks, suggesting a role even in procedural memory and emotional and cognitive functions (Habas, 2010).

Several studies have shown that the DNs can be altered in different neurological pathologies such as Friedreich’s ataxia (Solbach et al., 2014) and Alzheimer’s disease (Fukutani et al., 1999) where morphological changes within DNs have been detected. Subjects with Temporal Lobe Epilepsy (TLE) have shown less intuitive findings. In human, there are general reports of cerebellar atrophy in TLE patients (Hermann et al., 2005), while animal models have shown a direct involvement of the DNs: in particular, an experimental study on cats (Babb et al., 1974) showed that electrical stimulation of the DNs shortened and reduced the onset of seizures in various epilepsy models. In addition, from other studies investigating mouse models of epilepsy (Krook-Magnuson et al., 2014)(Kros et al., 2015) it emerged that neuro-stimulation of the DNs has greater effects in the inhibition of epileptic seizures than neurostimulation of the cerebellar cortex. While understanding the role of the DNs in epilepsy is beyond the scope of this work, it is important to indicate possible future applications of automatic DNs segmentation.

T1-weithed (T1-w) images are considered structural scans and generally this sequence is used for segmenting different brain regions. The DNs, unfortunately, do not show contrast on T1-w scans, while they are visible on T2-weighted (T2-w) images (Diedrichsen, 2006). This limitation might be one of the reasons why the involvement of DNs in human pathologies have been to date understudied. Currently, manual segmentation is still considered the gold standard for DNs segmentation (Acosta-Cabronero et al., 2017) (Lindig et al., 2019) (Akram et al., 2018) (Deoni and Catani, 2007), but it is time-consuming and suffers from inter- and intra-rater variability. A fully automatic segmentation is therefore desirable (Despotović et al., 2015). There is a published pilot study in 4 subjects (Ye et al., 2012) that proposes a fully automatic method to segment the DNs using DWI, which needs information obtained from tractography (requiring significant time). Another piece of work (Bermudez Noguera et al., 2019), for which only the abstract is available, proposes a deep learning approach to segment the DNs using as input multiple data including T1-w, T2-w images and Fractional Anisotropy (FA) maps. Using FA, or other quantitative map from DWI, to segment the DNs, though, precludes one to report the DNs’ metric (i.e. FA) as this would introduce a circular bias.

In reference (Bazin et al., 2018) the authors propose a fusion technique based on explicit shape modelling for the segmentation of CNs, starting from high-resolution 7T quantitative susceptibility mapping (QSM) of the cerebellum to accurately segment the DNs. In a recent piece of work (Li et al., 2019) a multi-atlas method was developed at 3T to segment iron-rich deep grey matter nuclei (including the DNs). This method, however, required QSM input images, not standardly acquired in clinical settings, and produced data not inherently registered with possible EPI-based quantitative sequences including DWI.

The purpose of this study is to segment the DNs for microstructure quantification of metrics acquired using the EPI readout as for DWI data. Indeed, it is possible to study the DNs using quantitative MRI methods that reflect micro-structural properties of tissues, such as those extracted from diffusion weighted imaging (DWI). In order to do so, segmentation masks of the DNs can be used to extract average values of quantitative metrics to be compared between populations of subjects (e.g. patients and healthy controls (HC)), to assess correlations with clinical scores or to monitor disease progression over time. Among the most interesting metrics there are quantitative parameters derived from clinically feasible Diffusion Tensor Imaging (DTI) or from advanced methods including Diffusion Kurtosis Imaging (DKI) (Jensen and Helpern, 2010), Neurite Density and Orientation Dispersion Imaging (NODDI) (Zhang et al., 2012), Composite Hindered And Restricted ModEl of Diffusion (CHARMED) (Assaf and Basser, 2005) and soma and neurite density imaging (SANDI) (Palombo et al., 2019). Given the typical resolution of DWI scans at 3T (2×2×2 mm^3^) and the low number of voxels included in segmentation masks of small structures such as the DNs, it is highly desirable to reduce the data manipulation due to post-processing steps (e.g. registration) and to have region segmented directly on DWI-space.

Moreover, it is essential that any automatic method is applicable with good performance to images of different quality and acquired with different scanners.

In the present study we developed a method to automatically segment DNs from non-diffusion weighted (b0) images, acquired as part of DWI scans. We specifically investigated three different approaches using high-resolution data derived from the Human Connectome Project (Essen et al., 2012): 1) atlas registration; 2) patch-matching; 3) a deep learning network-based method. The automatic segmentation masks obtained with each of these three methods were compared to the gold standard manual segmentation of DNs. The three automatic methods were subsequently tested in a second dataset of subjects involved in a TLE study. The resulting best approach was employed to compare DN volumes and average values of DWI metrics between patients and HC, in view of future clinical studies.

## 2 Methods

### 2.1 Subjects

#### HCP dataset

Pre-processed images of 100 healthy subjects scanned for the Human Connectome Project (HCP) were downloaded from the Connectome DB (http://db.humanconnectome.org) (Van Essen et al., 2013). 24 of these subjects were discarded because of severe cerebellar artefacts. The remaining 76 subjects (43 Females, 29.41±3.62 years) were analysed and used to develop an automatic DNs segmentation method.

#### TLE dataset

A second dataset was used to test the performance of the three automatic segmentation methods and to pilot its clinical applicability. 84 subjects were recruited for an Italian multi-centre research project on TLE. Subjects were divided in three groups: 34 HC (16 Females, 31.97±7.73 years), 21 patients with left TLE (LTLE; 13 Females, 33.29±11.68 years) and 29 patients with right TLE (RTLE; 17 Females, 37.97±9.86 years). For each subject handedness was recorded, as reported in Table1.

**Table 1:** Handedness (left; right) of the subjects involved in the TLE study.

| Groups | Left | Right |
| --- | --- | --- |
| HC | 6 | 20 |
| LTLE | 1 | 19 |
| RTLE | 4 | 25 |

### 2.2 MRI protocol

#### HCP dataset

MR images were acquired using a customised Siemens 3T Connectome Skyra scanner with a dedicated gradient insert (diffusion: Gmax = 100 mT/m, max slew rate = 91 mT/m/ms; readout/imaging: Gmax = 42 mT/m, max slew rate = 200 mT/m/ms), a 32-channel receive head coil and standard shim coils (WU - Minn Consortium Human Connectome Project, 2017). We downloaded DWI data with minimal pre-processing, co-registered with T1-w data at a resolution of 1.25×1.25×1.25 mm^3^ and matrix size of 145×174×145 (WU - Minn Consortium Human Connectome Project, 2017). The DWI acquisition included 18 volumes with b=0 s/mm^2^.

#### TLE dataset

MR images were acquired using a Siemens 3T MAGNETOM Skyra scanner with standard gradients and a 32-channel receive coil.

##### DWI

spin-echo EPI with TR=8400 ms, TE =93 ms, 90 volumes with b-value=1000/2000 s/mm^2^ (45 different diffusion weighted gradient directions per b-value) and 9 volumes with b=0 s/mm^2^. The spatial resolution was 2.24×2.24×2 mm^3^, with a field of view of 224×224 mm^2^ and matrix size of 100×100×66 voxels.

##### T1-w

high-resolution 3D T1-w (T1w) volume acquired with a multi-echo FLASH sequence: TR=19 ms, six equidistant TE from 2.46 to 14.76 ms, flip angle 23°. The spatial resolution was 1×1×1 mm^3^, with a field of view of 256×232×176 mm^3^.

### 2.3 DWI processing

For both datasets we computed the mean of the b0 volumes belonging to a single subject, from now on referred to as 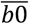. Moreover, for the TLE subjects, the following quantitative metrics were extracted using DESIGNER (https://github.com/NYU-DiffusionMRI/DESIGNER) (Ades-Aron et al., 2018): Axial Diffusivity (AD), Radial Diffusivity (RD), Mean Diffusivity (MD) and FA from DTI fitting (Alexander et al., 2007) and Axial Kurtosis (AK), Radial Kurtosis (RK) and Mean Kurtosis (MK) from DKI fitting (Jensen and Helpern, 2010).

### 2.4 DNs segmentation

To develop an automatic DNs segmentation method, we used the average 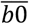 images of each HCP subject. Manual segmentation was used as ground truth (GT). Automatic DN masks, from the three different automatic segmentation methods, were then compared to the GT masks. The automatic methods were subsequently applied to a clinical context, i.e. the TLE dataset. Quantitative evaluation of each method’s performance against the GT was carried out by calculating three scores: the Dice Similarity Coefficient (DSC), the True Positive Rate (TPR) and the Positive Predictive Value (PPV) (see Quantitative Evaluation below, paragraph 2.6).

#### Ground Truth (GT) – manual segmentation

The 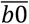 images of all HCP subjects were manually segmented by two raters: rater 1 and rater 2. These raters used two different software packages: rater 1 used Mango (http://ric.uthscsa.edu/mango/mango.html) and rater 2 used Jim (http://www.xinapse.com/j-im-8-software). The average inter-rater variability was evaluated first by calculating the DSC between the two manual segmentation masks from raters 1 and 2 for each HCP subject and then by averaging the DSCs of all 76 HCP subjects. Moreover, 6 subjects were segmented twice by the same operator (rater 1) on different days to calculate the intra-rater variability as the average DSC between the two manual segmentation masks for rater 1. We arbitrary choose rater 1 segmentations for training. For the TLE dataset, rater 1 manually segmented the 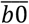 of 18 subjects (6 for each group) in order to have a GT (GT_TLE_) for this independent dataset.

#### Atlas-based method: SUIT

The toolbox SUIT (A spatially unbiased atlas template of the cerebellum and brainstem) (diedrichsenlab.org/imaging/suit.htm) is an open source extension of SPM (Statistical Parametric Mapping, https://www.fil.ion.ucl.ac.uk/spm/) available for Matlab (The MathWorks, Inc., Natick, MA, United States of America). SUIT (Diedrichsen, 2006)(Diedrichsen et al., 2011) is an atlas-based method for cerebellar segmentation that performs a non-linear registration between a template (standard space) and the image to segment. The resulting transformation is then applied to an atlas defined in the standard space and its labels are warped into the subject space. The SUIT toolbox provides an anatomical atlas of the cerebellum that includes the CNs, hence also the DNs. SUIT requires the user to first register the T1w images of each subject to the template; the inverse transformation is then used to warp DN labels from standard-space to subject-space. As the T1w images of the HCP dataset are already co-registered with the respective DWI, the DN segmentations obtained with SUIT are already in DWI space.

#### Pre-processing (OPAL and CNN)

In order to segment DNs with OPAL and CNN we applied two pre-processing steps:

1. Intensity normalization: The mean intensity and its standard deviation were calculated for each subject’s 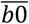 volume, considering the intensity value only of voxels belonging to the brain. Each 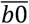 was normalized in order to obtain zero mean and standard deviation equal to 1 for all subjects.
2. Crop: In order to reduce the computational time, images were cropped around the cerebellum reducing the size of axial slices from 145×174 to 86×71 voxels, centered at the (73, 45) voxel in-plane co-ordinate position.

#### Patch-matching method: OPAL

OPAL (Optimized Patch Match for Label fusion) (Giraud et al., 2016) joins information from different templates to obtain the desired segmentation. OPAL is an evolution of the Patch Match algorithm (Barnes et al., 2009), implemented in C++ (https://github.com/KCL-BMEIS/NiftySeg/).

We built up a database of 46 subjects providing the following information for each subject: 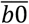 images, the corresponding masks of the cerebellum and the DN GTs. This database was intended as a collection of reference templates. The DNs segmentation of each new subject was performed by dividing images into patches and comparing each patch with those from the reference templates, looking for the most locally similar match. The output is a probabilistic map of the DNs. We divided the 30 subjects into two equal sets, one for validation and one for testing. We used the validation set to select the probability thresholds (0.1, 0.2, 0.3, 0.4, 0.5) for binarizing the DN masks, where a lower threshold corresponds to larger DN masks. For each threshold and for each validation subject we calculated the DSC between the DN masks and the GTs. We selected the threshold that maximised the mean DSC and we assessed the performance of OPAL on the remaining 15 test subjects for an unbiased performance estimate.

#### Deep-learning method: CNN

A CNN (Convolutional Neural Network) was implemented with Matlab19a using the available Deep Learning Toolbox.

##### CNN architecture

The architecture used here was inspired to the one used for segmenting the spinal cord grey matter (Perone et al., 2018). This architecture was based on dilated convolutions and on removal of pooling layers, responsible for information loss. This type of convolution expanded receptive fields without increasing the number of parameters (Khan et al., 2018). The network implemented required as input a two-dimensional (2D) image, oriented in the axial plane. The architecture is shown in Figure 1.

**Figure 1:**
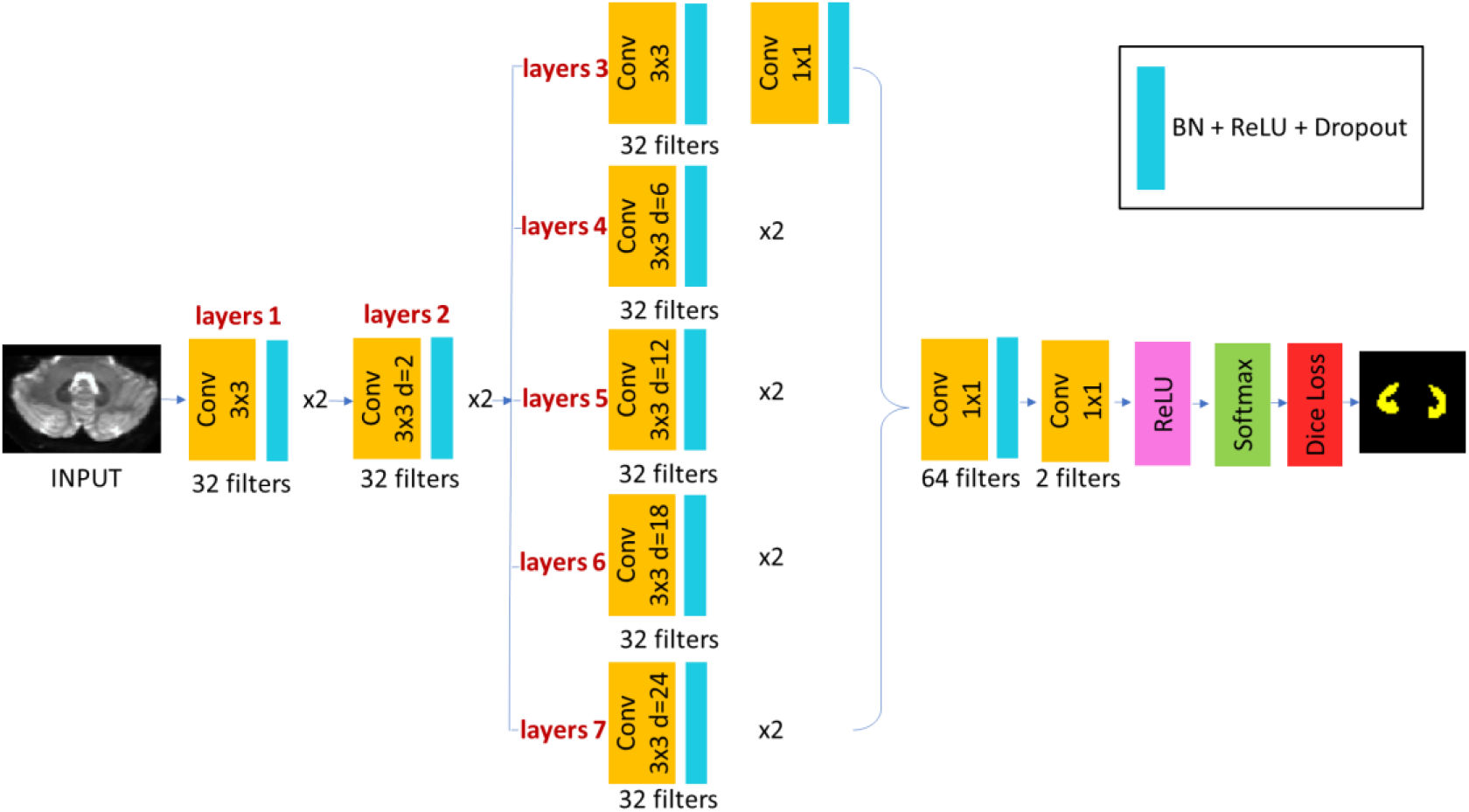
Scheme of the CNN architecture adopted here.

All convolutional layers have a zero-padding of type “same” (Dumoulin and Visin, 2016). Therefore, the dimensions of each layer’s output do not differ from those of the layer’s input. For each layer the neurons are activated by the ReLU (Rectifier Linear Unit) function (Aylward et al., 2017).

The architecture of the CNN is the following:

- Input layer (INPUT) receiving the input images and treating each voxel as a neuron of the input layer;
- Two layers of standard convolution (layers 1);
- Two layers of dilated convolution with dilatation factor d = 2 (layers 2);
- Five branches in parallel, each branch with two layers of convolution:

- In the first branch there is a standard convolution: for the first layer the kernel dimension is 3×3 while for the second layer it is 1×1 (layers 3);
- In the second branch there is a dilated convolution with d=6 (layers 4);
- In the third branch there is a dilated convolution with d=12 (layers 5);
- In the fourth branch there is a dilated convolution with d=18 (layers 6);
- In the fifth branch there is a dilated convolution with d=24 (layers 7).

Each output of these parallel branches is concatenated in the third dimension and followed to:

- A convolution layer that uses 64 filters of dimensions 1×1;
- A convolution layer that uses 2 filters of dimensions 1×1;
- A Softmax layer (Aylward et al., 2017) that represents the activation function for classification;
- A Loss layer.

The convolutional layers have 32 filters with dimension 3×3 except for the second layer of layers 3, which is 1×1, and the last two layers. Except for the last 1×1 convolution, each convolution layer is followed by batch normalization (Ioffe and Szegedy, 2015) and dropout (Srivastava et al., 2014) (Khan et al., 2018).

Due to the imbalance between the class of belonging to the DN and the non-belonging class (i.e. background), we decided to use the Dice Loss as loss function, based on the DSC and robust to class imbalance (Fidon et al., 2018). We used the Adam optimizer (Kingma and Ba, 2017) with a small learning rate of η=0.001 for setting the weights of the CNN parameters.

##### Training

In order to reduce overfitting, a commonly used technique known as data augmentation was applied, increasing the size of the training dataset. Four different transformations were considered: rotation, translation, scaling and elastic deformation. These transformations were applied to input 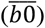 - desired output (GT) pairs. Data augmentation was applied independently on each slice with a probability of 0.5 for each transformation. The parameters used are reported in Table2.

**Table 2:** Range of parameters used for the transformations applied during the data augmentation step of the CNN optimisation. For each slice, with 0.5 probability, a random number within this range was assigned to each transformation. For elastic deformation α represents the scale factor, while σ represents the standard deviation of the Gaussian filter.

| Transformation | Parameter range |
| --- | --- |
| Rotation | $[-4.6^\circ, 4.6^\circ]$ |
| Shift | $[-3, 3]$ in x and y direction |
| Scaling | $[0.98, 1.02]$ with bicubic interpolation |
| Elastic Deformation | $\alpha = 4$ and $\sigma = 30$ |

The original 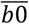 images plus those from data augmentation and the corresponding GT masks were provided as input to the CNN for training. To speed up training, however, only slices containing the DN (on average 8 per subject) were included as selected from the GT masks. The hyperparameters that must be chosen a priori before training were the batch size, the dropout and the number of epochs. For tuning these hyperparameters we tried a number of combinations (45 in total), as reported in Table3.

**Table 3:**
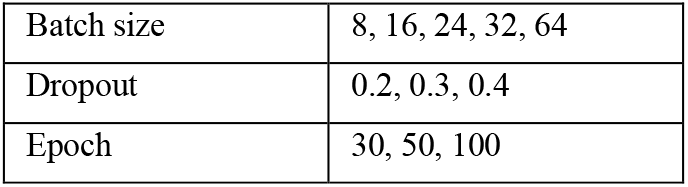
Hyperparameters values combined in 45 different hyperparameters sets.

|  |  |
| --- | --- |
| Batch size | 8, 16, 24, 32, 64 |
| Dropout | 0.2, 0.3, 0.4 |
| Epoch | 30, 50, 100 |

For each combination of hyperparameters, a Monte Carlo 10-folds cross validation was performed as follows: firstly, we randomly extracted 6 of the 76 subjects to be used as a test set. Then, the remaining 70 subjects were randomly split into 60 subjects for training and 10 subjects for validation; this step was repeated for each of the 10 folds. The Monte Carlo 10-folds cross validation randomly selects subjects for the training and the validation set, therefore it is possible that a subject is never included or can be used more than once in the validation set.

The steps used for CNN training are shown in Figure 2: 1) for each fold of each combination of hyperparameters we calculated the DSC for the subjects included in the validation set (10 subjects); 2) we calculated the mean DSC for each hyperparameters combination by averaging the DSCs of the 10 folds; 3) we chose the combination of hyperparameters that maximized the average DSC; 4) among the 10 CNN that were trained with the best hyperparameters combination, we chose the one with the maximum DSC. Set the hyperparameters, we used the 6 test subjects for an unbiased estimate of the CNN performance.

**Figure 2:**
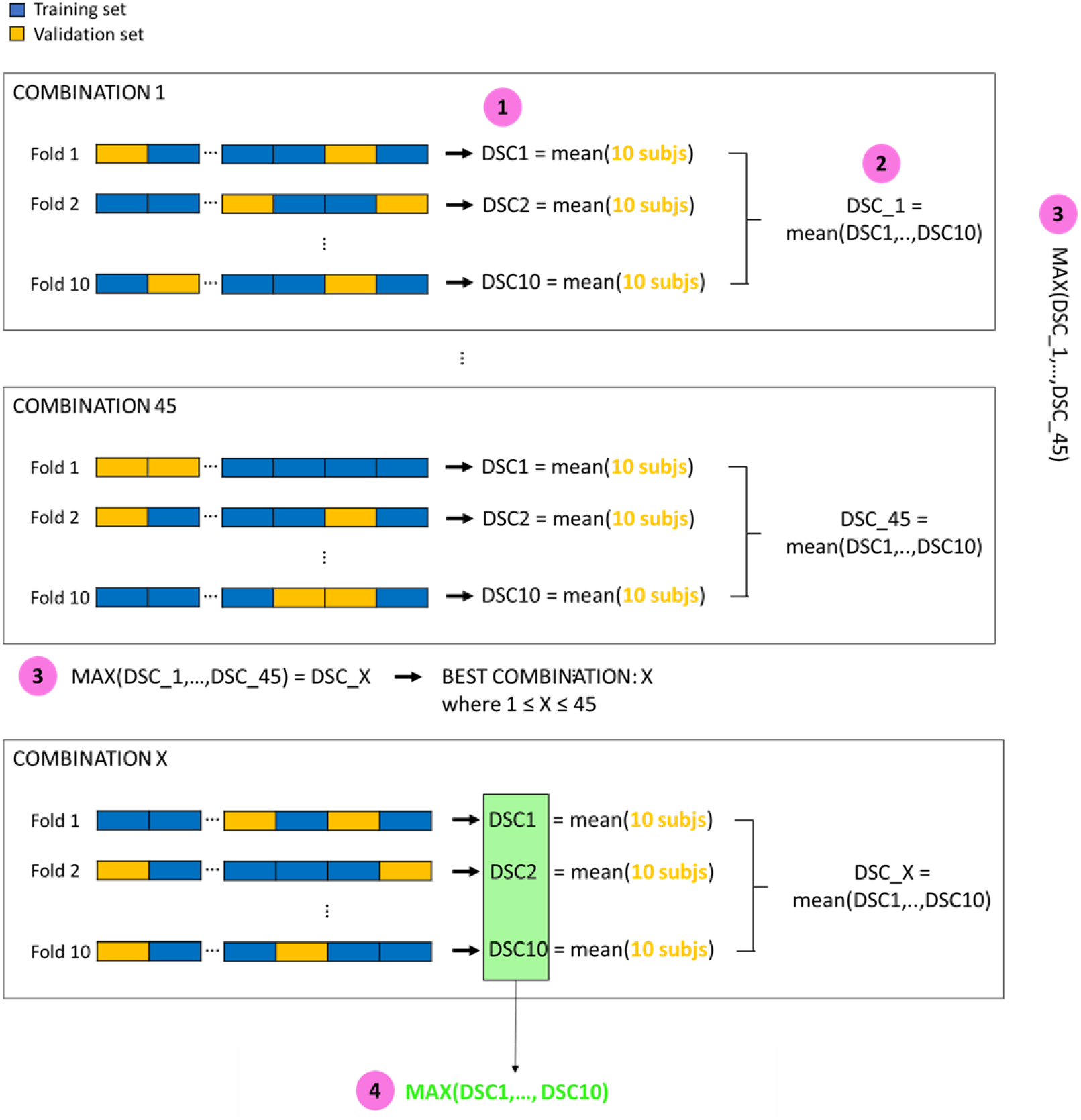
Steps followed for hyperparameters optimization and CNN training.

### 2.5 Post processing for OPAL and CNN

Both OPAL and CNN labeling identified a number of false positive (FP) voxels as belonging to the DNs located in different brain regions, sometimes very distant from the DNs themselves. In order to remove these FP voxels, an automated post processing step was implemented: the DN masks obtained with SUIT were dilated twice (fslmaths, FMRIB Software Library (FSL), https://fsl.fmrib.ox.ac.uk/fsl/fslwiki) and used to mask the DN masks generated by OPAL and CNN.

### 2.6 Quantitative evaluation

For each method, performance was tested by comparing automatic DNs against GT masks using three different scores(Prados et al., 2017).

- Dice Similarity Coefficient (DSC), i.e. the overlap between two binary masks:

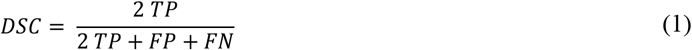

where TP indicates True Positive and FN False Negative. DSC ranges [0-1].
- Sensitivity or True Positive Rate (TPR):

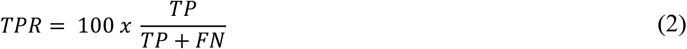

TPR ranges [0-100] with low TPR indicating a bias towards under-segmentation.
- Precision or Positive Predictive Value (PPV):

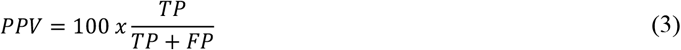

PPV ranges [0-100] with low PPV indicating a bias towards over-segmentation

Specificity or True Negative Rate (TNR) was not considered because the two classes (DN and background) are unbalanced, causing high and non-informative TNR values.

### 2.7 Comparison of automatic methods

We calculated DSC, TPR and PPV for each automated method. For OPAL and CNN we calculated these scores, on the validation and test sets, before and after post processing. Since SUIT is an atlas-based method we calculated these scores on the whole dataset, while for OPAL we exclueded the 46 subjects used as template. Regarding CNN, the scores were calculated for the validation (10 subjects) and test (6 subjects) sets for each of the 10 folds corresponding to the optimal set of hyperparameters. For each method we calculated the group average of these scores.

For the CNN we calculated two average values: the first one by averaging between the 10 folds corresponding to the best combination of hyperparameters, while the second one by averaging only results obtained with the network chosen as the final CNN (the one with the best perfomance) among the 10 networks.

### 2.8 Clinical application to TLE data

Figure 3 shows the pipeline used to segment the DNs on the independent TLE dataset.

**Figure 3:**
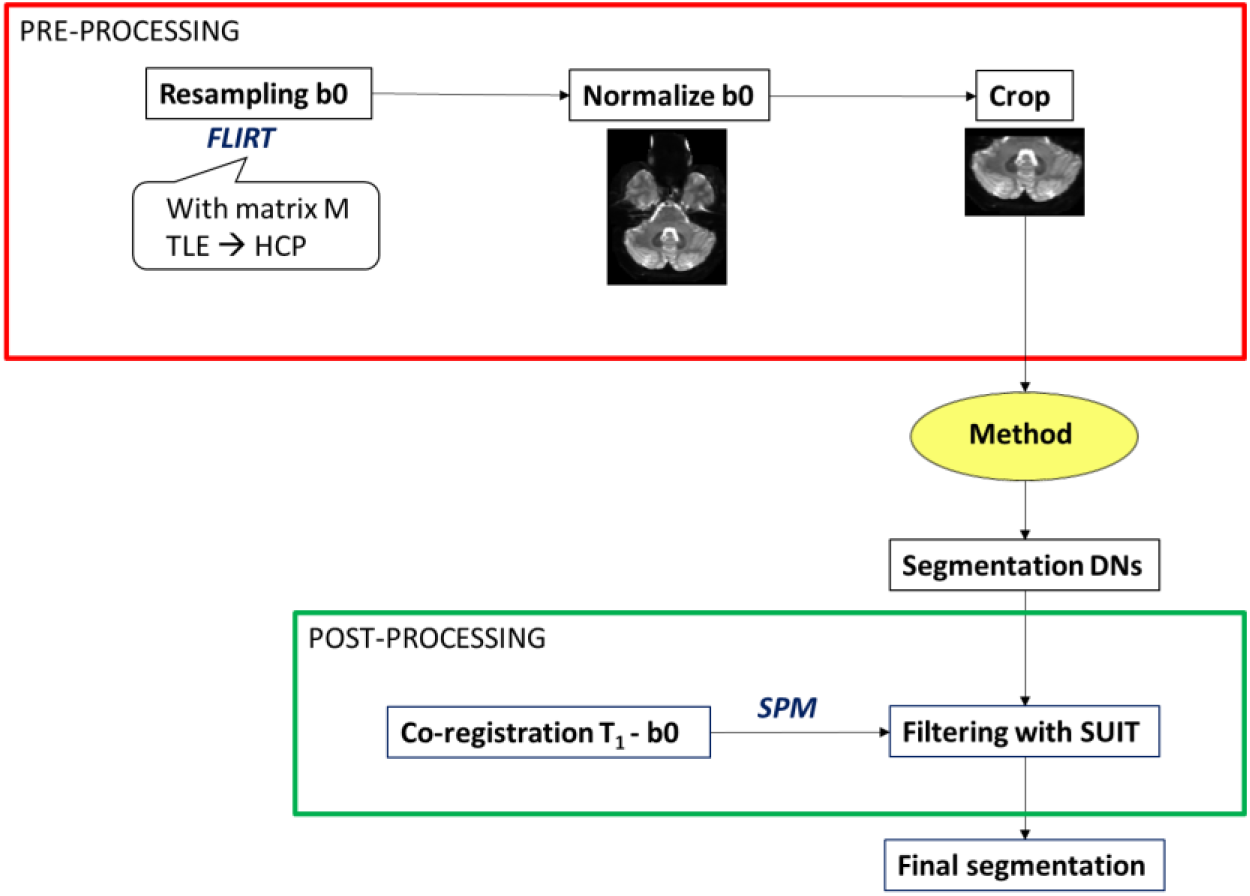
Pipeline followed to segment TLE 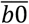 images.

#### TLE data pre-processing and DNs segmentation

The spatial resolution of the TLE 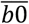 images was lower than that of the HCP dataset, so TLE 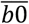 images were resampled to match the HCP resolution using FSL FLIRT (FMRIB’s Linear Image Registration Tool) before applying each segmentation method. To remove the FPs, T1w images were registered to 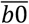, using a rigid registration in SPM. The resulting DN masks were resampled to their original spatial resolution for quantitative analysis of parameter maps by applying the inverse of the roto-translation matrix. GT_TLE_ segmentations were used to assess performance of the three methods.

We selected the best automatic DNs segmentation method based on the performance on both the HCP dataset and on the 18 TLE subjects. The best method was applied to all TLE subjects in order to extract quantitative DNs parameters from DWI.

#### DN structural and microstructural characteristics in TLE patients

Quantitative measures of each DN (right and left DN independently) were extracted to perform statistical comparisons between groups of TLE patients and HC. These measures were: 1) the volume of each DN; 2) the average value of DTI metrics (AD, RD, MD and FA) for each DN; 3) the average value of DKI metrics (AK, RK and MK) for each DN. Lateralization of volumes and metrics between right and left values was investigated using an Asymmetry Index (AI) (Bonekamp et al., 2007):

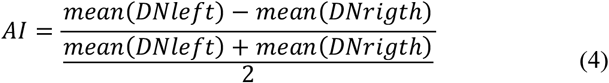

The range of AI values is [-2; 2] where 0 indicates perfect symmetry.

We considered a total of 24 measures for each subjects. Statistically significant differences of these measures between the three groups were investigated using SPSS (IBM, Armonk, NY, United States of America) as exploratory work.

Age, gender and handedness were compared between groups and were taken into account for the final statistical comparison. A general linear model (GLM) univariate analysis was implemented using as covariates those variables not homogeneous between groups. 24 GLM univariate comparisons, with α=5%, were performed to explore which variables could significantly differentiate the three groups. Subsequently GLM univariate analysis was repeated for each metric in pairwise group comparisons.

## 3 Results

The inter-rater variability of the manual segmentations resulted in a DSC = 0.8066 ± 0.0575. Intra-rater variability produced a DSC = 0.7927 ± 0.0369. In Figure 4 DN masks of a randomly selected subject are displayed, where each method (red) is compared to the GT (yellow). OPAL probability threshold was set to 0.4. The Monte Carlo 10-folds cross validation of the CNN provided the best results with this set of hyperparameters: batch size = 24, dropout = 0.2 and number of epochs = 100.

**Figure 4:**
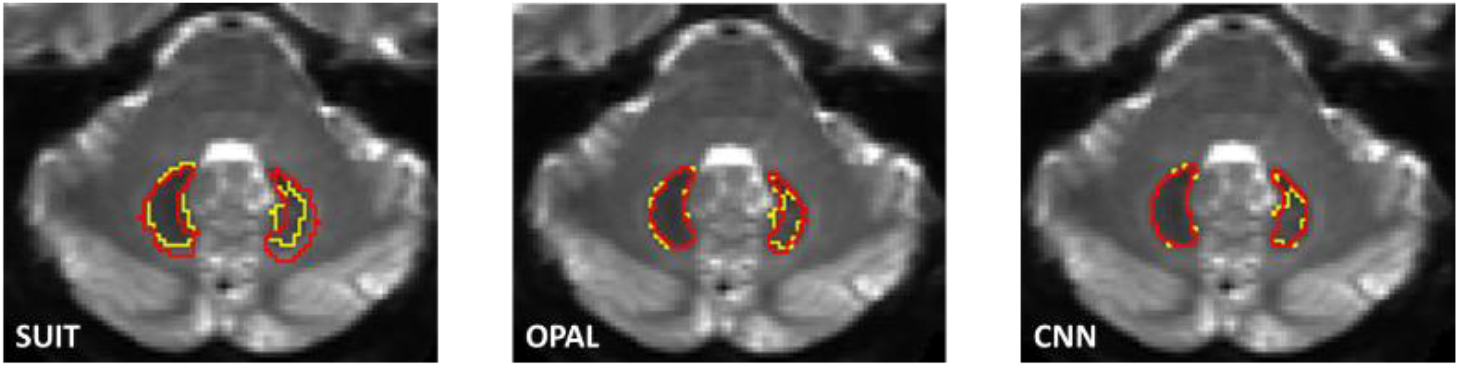
Segmentation masks obtained with the different methods for a randomly selected subject (SUIT, OPAL and CNN). Each image shows the overlap of the segmentation obtained with the respective automated method (red) overlaid with the GT (yellow).

### 3.1 Comparison of the three automatic methods

DSC, TPR and PPV scores for SUIT (mean ± standard deviation) are DSC = (0.4907 ±.0.0793); TPR = (86.3444 ±6.6154) and PPV = (34.9475 ±7.6264). Table 4 reports DSC, TPR and PPV scores for OPAL and CNN for both the validation set and the test set. For CNN are reported both the average score from the 10 networks obtained during the 10-folds cross validation that resulted in the chosen hyperparameters and the scores obtained with the final CNN network set as the best performer amongst the same 10. Scores after the post-processing step are reported with DSCs shown also without the post processing step (in brackets) for comparison.

**Table 4:**
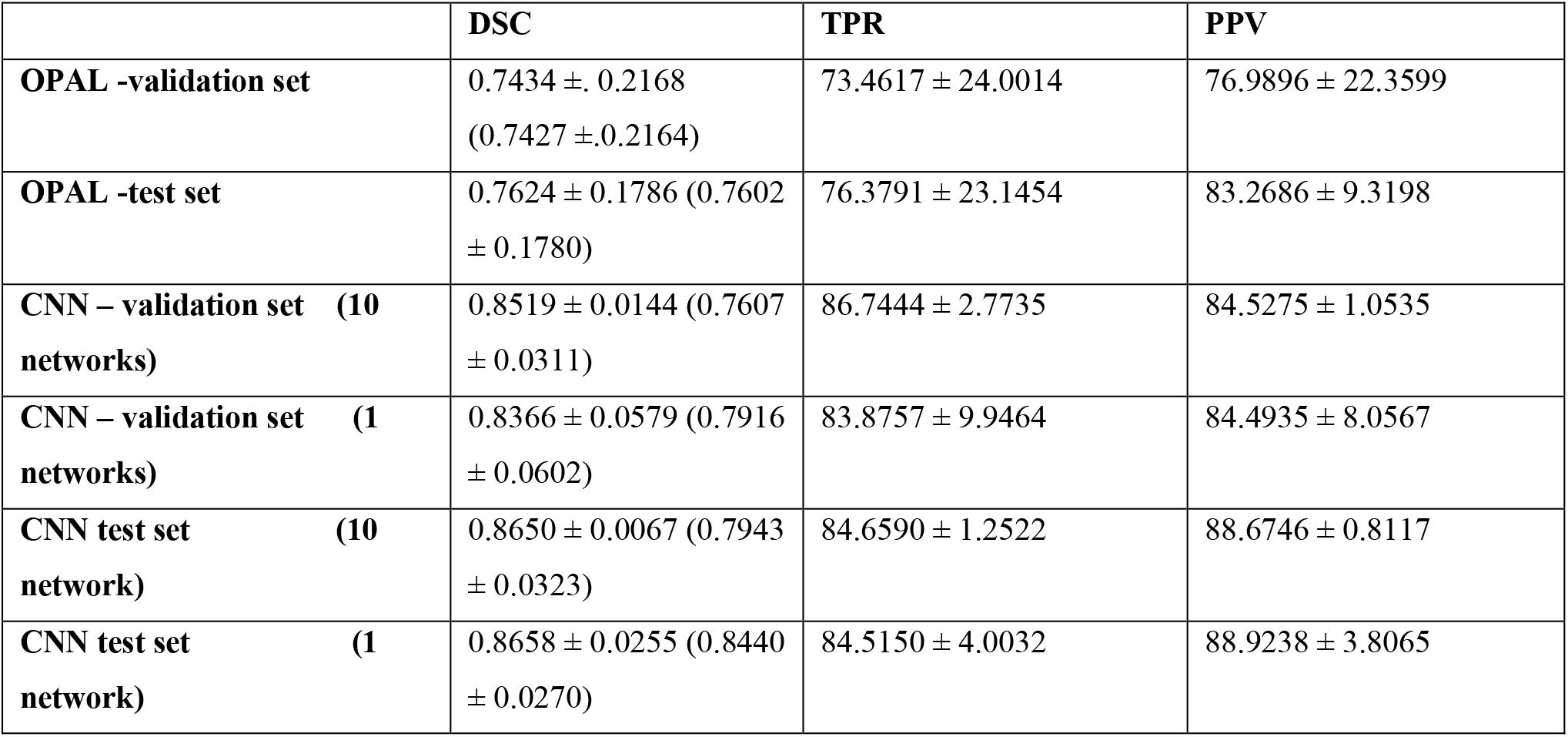
OPAL and CNN performance after the post processing step. For CNN, two sets of scores are reported: 1) Average scores from the 10 networks with the chosen hyperparameters; 2) metrics from results obtained with the CNN network chosen as the best performer. For DSC, in bracket we reported the values before the post processing step to remove false positives.

The best performance was achieved by CNN (DSC = 0.8658 ± 0.0255). The CNN scores were followed by OPAL (DSC = 0.7624 ± 0.1786). SUIT performed worst, thus producing the lowest scores (DSC = 0.4907 ± 0.0793).

### 3.2 Application to TLE dataset

Table 5 reports DSC, TPR and PPV scores between GT_TLE_ and the segmentation obtained with each automatic method. For OPAL it was necessary to reset the probability threshold to 0 as 0.4 (set for the HCP data) eliminated true positives. Overall scores were: DSC = 0.1322 ± 0.1512, TPR = 7.7931 ± 9.2878 and PPV = 55.2716 ± 50.8794. CNN outperformed the other methods with a DSC = 0.7368 ± 0.0799.

**Table 5:** Comparison of SUIT, OPAL and CNN against GT on 18 TLE subjects.

|  | DSC | TPR | PPV |
| --- | --- | --- | --- |
| **SUIT** | $0.4145 \pm 0.1023$ | $84.3647 \pm 8.4051$ | $27.9597 \pm 8.6905$ |
| **OPAL** | $0.4522 \pm 0.1178$ | $84.3277 \pm 16.0649$ | $28.6451 \pm 12.1937$ |
| **CNN** | $0.7368 \pm 0.0799$ | $88.6787 \pm 4.5745$ | $65.7410 \pm 10.6841$ |

From the statistical tests run to compare the TLE sub-groups, it resulted that age was not homogeneous in the three groups (p-value = 0.017), while gender was matched (p-value = 0.491) and handedness was balanced (p-value = 0.301). Therefore, the statistical comparisons of DWI metrics included age as a GLM covariate.

We found some significant differences between the three groups: AD of the left DN (p-value= 0.024), MD of the left DN (p-value = 0.039) and volume of the right DN (p-value = 0.014). Figure 5 shows boxplots of these metrics for each group. Pairwise comparisons between two of the three groups showed that: AD of the left DN is significantly different between LTLE and RTLE patients (p-value = 0.004), MD of the left DN is significantly different between LTLE and RTLE patients (p-value = 0.016), the volume of the right DN is significantly different between HC and LTLE patients (p-value = 0.049) and between HC and RTLE patients (p-value = 0.010). Moreover from pairwise comparisons other metrics resulted significant differences: volume of the left DN is significantly different between HC and RTLE patients (p-value = 0.027) and RD of the left DN is significantly different between HC and LTLE patients (p-value = 0.044). In Figure 6 are reported the boxplots of these metrics for each group.

**Figure 5:**
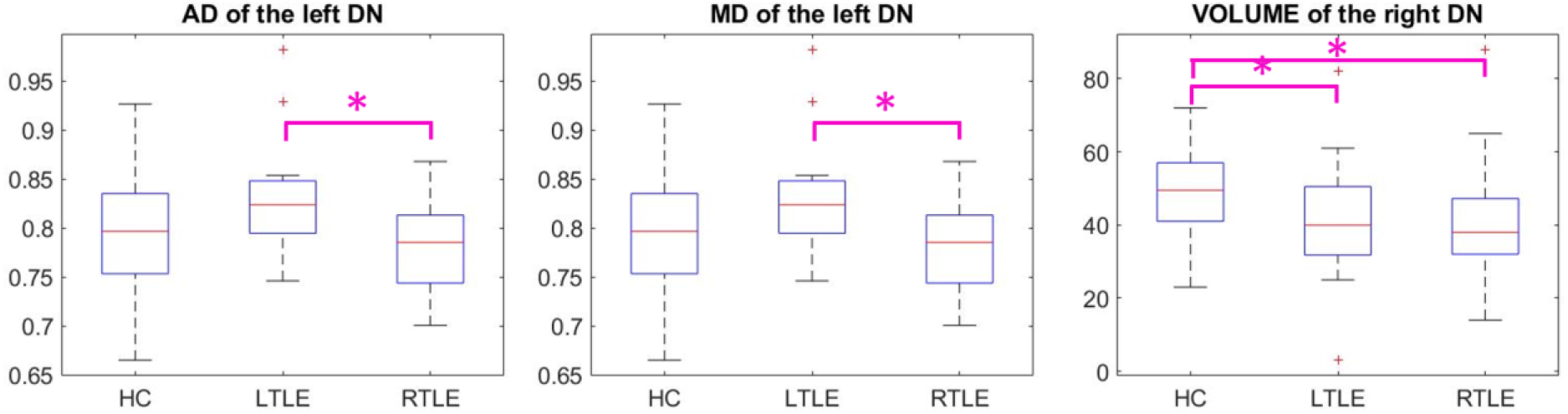
Boxplots of the measures that resulted statistically different (p<0.05) between the three groups of HC, LTLE and RTLE patients: AD of left DN, MD of the left DN and volume of right DN. Pairwise comparisons that resulted significantly different are highlighted with an asterisk.

**Figure 6:**
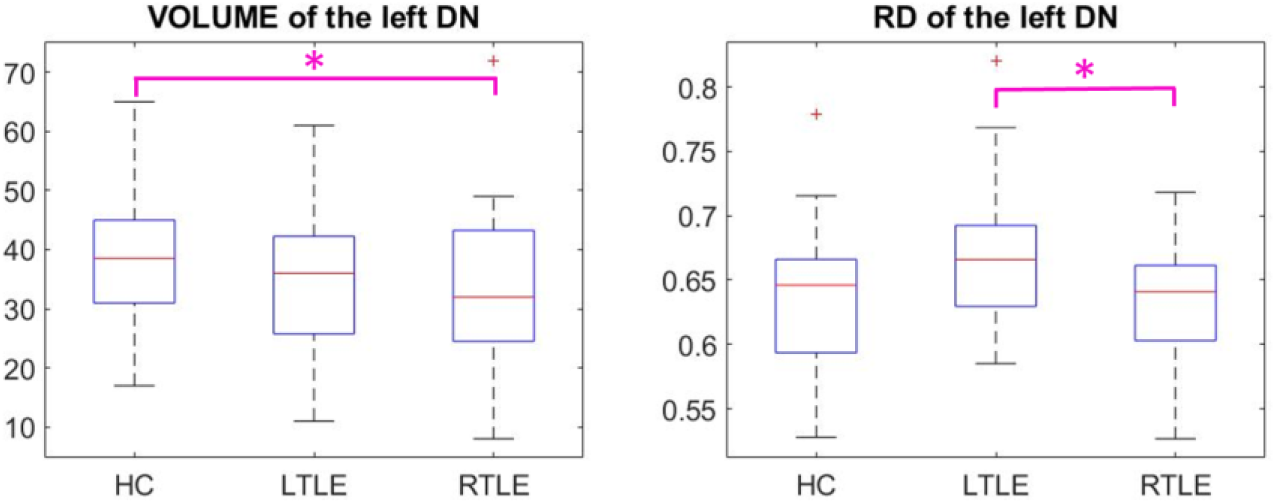
Boxplots of the measures that resulted statistically different (p<0.05) from pairwise comparisons (highlighted with an asterisk): volume of the left DN and RD of the left DN.

## 4 Discussion

In this work we proposed an automatic DNs segmentation method that uses non-DW 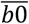 images from a DWI dataset. Specifically, analysis of DSC scores highlighted performances comparable with inter- and intra-raters segmentation (DSC > 0.7). The use of 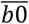 images, inherently co-registered with DWI data, instead of high resolution T1w structural scans, allows the user to apply the masks directly to microstructural parameter maps obtained for clinical research studies.

On HCP data, segmentation masks obtained with OPAL and CNN were more accurate than the over-segmented DNs obtained with SUIT. Furthermore, DSC, TPR and PPV average values were all superior for segmentations using CNN compared to OPAL.

OPAL applied to TLE data had a much worse performance (even after changing the threshold). This indicates that OPAL, which here used a reference database constructed using HCP data, cannot segment images acquired on a different scanner and with a worse resolution. Possibly, to improve the performance of OPAL, one would need to build a more appropriate database of reference templates.

Therefore, the implemented CNN outperforms OPAL and can be considered the best automated segmentation method of DWI images among the ones tested here (the code for the CNN is publicly available at https://github.com/marta-gaviraghi/segmentDN/).

One further major advantage of CNN over OPAL lies in its greater transferability across sites and users. Indeed, OPAL requires that the database of 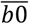 s and associated GTs is available to segment the DNs of new subjects. Conversely, CNN needs a database of images and GTs only for training, but after the network has learnt the association between images and segmentations, the reference images are no longer needed. One could question also the dependency of the method on the geometrical acquisition parameters, but here we demonstrated that the method worked well (DSC>0.73) also on a completely different dataset, acquired on a standard clinical 3T scanner and with a much coarser voxel resolution. We recommend that the performance of the CNN is assessed on a subset of images before systematically applying it to a new DWI datasets.

The CNN was applied to the 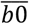 data of the TLE dataset to segment the DNs and study their microstructural properties in a group of patients affected by TLE. While understanding the DNs involvement in TLE requires a dedicated study comparing regions from the entire brain (and not just the DNs), it was very interesting to see that the DN masks obtained from the 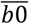 images could be easily applied to DTI and DKI metrics and be used for some very preliminary assessment. The statistical comparison showed that the right DN volume is reduced in both RTLE and LTLE with respect to HC. The volume reduction of the right DN in TLE patients could indicate atrophy of this cerebellar nucleus, but to understand the source of such alteration one should also consider what happens to the underlying microstructure and hence assess parameters from, for example, DTI or DKI fitting of the data as it was performed here. From our exploratory comparisons, AD and MD seem to be the most affected metrics, which might simply relate to a different proportion of white and grey matter structures captured by the masks in different groups. To disentangle the source of such changes, though, future studies should consider advanced microstructural models that probe more specific biophysical properties such as neuronal density, orientation dispersion and soma compartments (Zhang et al., 2012)(Palombo et al., 2019). These preliminary results support the hypothesis that DNs might be involved in TLE, consistently with previous studies in animal models of epilepsy (Babb et al., 1974) (Krook-Magnuson et al., 2014) (Kros et al., 2015). The extent of such involvement must be explored further within a dedicated clinical study that correlates DN alterations with that of other brain regions, considering also clinical/anamnestic data such as comorbidities and treatment (Mavroudis et al., 2013).

Methodologically, given the coarse resolution of DWI data, a potential limitation of using 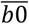 images is that it is not possible to extract the convoluted surface of the DNs and to specifically extract their grey matter. Current structural scans used for the segmentation of small regions, i.e. 3D T1-w scans, do not show contrast in the CN areas. If a detailed reconstruction of the DNs shape and size is considered a fundamental aspect for a specific study, a dedicated sequence with optimized contrast (e.g. based on T2 or T2* properties or QSM) and image resolution (e.g. to achieve sub-millimetre voxel size) should be considered, at the expense of longer acquisition times. For the purpose of our study, 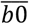 images served the purpose of achieving a significant improvement over the SUIT segmentation without resorting to additional MR sequences and longer acquisition time. Furthermore, the demonstrated translation of the CNN from the HCP to a clinical scanner DWI data is very encouraging and makes this CNN possibly viable for other applications that use EPI-readouts; future work could therefore investigate transability of the proposed CNN to study functional MRI activations of the DNs in relation to their microstructure characteristics.

## 5. Conclusion

We proposed an automatic segmentation of the DNs using a fully automated method. The CNN implemented here can segment images with a spatial resolution and acquisition protocol different from the training set. By using the proposed CNN on a cohort of subjects affected by TLE we detected asymmetric microstructural changes within the DNs, which should be further investigated in dedicated studies. Future work could consider multimodal datasets including as input images with different MRI contrasts and an expanded GT database for training.

## Acknowledgements

Data were provided by Human Connectome Project, WU-Minn Consortium (Principal Investigators: David Van Essen and Kamil Ugurbil; 1U54MH091657) funded by the 16 NIH Institutes and Centers that support the NIH Blueprint for Neuroscience Research; and by the McDonnell Center for Systems Neuroscience, Washington University. 3TLE is a multicentric research project granted by Italian Health Ministry (NET2013-02355313): Magnetic resonance imaging in drug-refractory temporal lobe epilepsy: standardization of advanced structural and functional protocols at 3T, to identify hippocampal and extra-hippocampal abnormalities. An acknowledgment for patients recruitment within this project to Carlo Andrea Galimberti. Acknowledgments to the UCL-UCLH Biomedical Research Centre for ongoing funding; the European Union’s Horizon 2020 research and innovation programme under grant agreement No. 634541, Spinal Research (UK), Wings for Life (Austria), Craig H. Neilsen Foundation (USA) (jointly funding the INSPIRED study), Wings for Life (#169111), the UK Multiple Sclerosis Society (grants 892/08 and 77/2017).

